# Tau tubulin kinase 1 and 2 regulate ciliogenesis and human pluripotent stem cells–derived neural rosettes

**DOI:** 10.1101/2023.01.18.524283

**Authors:** Lucia Binó, Lukáš Čajánek

## Abstract

Primary cilia are key regulators of embryo development and tissue homeostasis. However, their mechanisms and functions, particularly in the context of human cells, are still unclear. Here, we analyzed the consequences of primary cilia modulation for human pluripotent stem cells (hPSCs) proliferation and differentiation. We show that neither activation of the cilia-associated Hedgehog signaling pathway nor ablation of primary cilia by CRISPR gene editing to knockout Tau Tubulin Kinase 2 (TTBK2), a crucial ciliogenesis regulator, affects the selfrenewal of hPSCs. In addition, we demonstrate that TTBK1, a closely related kinase without previous links to ciliogenesis, is upregulated during hPSCs-derived neural rosette differentiation to regulate primary cilia formation together with TTBK2. Finally, we show that TTBK1/2 and primary cilia are implicated in the regulation of the size of hPSCs-derived neural rosettes.

## Introduction

Cilia are hair-like organelles protruding from the surface of most cells. While the motile cilia are perhaps the oldest organelles ever known (Beales & Jackson, 2012), the function of the single non-motile primary cilium has for a long time been enigmatic. It is now appreciated that primary cilium contains receptors and effectors of several signaling pathways, such as the Hedgehog (HH) pathway (Anvarian *et al*, 2019; Ingham, 2022). Conversely, primary cilia govern important aspects of embryonic development and tissue homeostasis (Bangs & Anderson, 2017; Ingham, 2022). Defects in the assembly and function of primary cilia cause diseases such as Bardet-Biedel syndrome, Joubert syndrome, or nephronophthisis, collectively termed ciliopathies (Reiter & Leroux, 2017). Cilia status also affects certain types of cancer, such as medulloblastoma or basal cell carcinoma (Han & Alvarez-Buylla, 2010). Therefore, targeting cilium-related pathways may represent a propitious therapeutic strategy.

The fully grown primary cilium is composed of the mother centriole-derived basal body, the transition zone, and the microtubule-based axoneme enclosed within a ciliary membrane (Satir *et al*, 2010). The cilium assembly is initiated at the distal end of the mother centriole, by the coordinated action of distal appendage components (such as CEP83 and CEP164) and Tau Tubulin kinase 2 (TTBK2) (Goetz *et al*, 2012; Čajánek & Nigg, 2014; Tanos *et al*, 2013; Graser *et al*, 2007). TTBK2 kinase activity is essential for ciliogenesis - no cilia are formed in cells devoid of the active kinase. Following the delivery and docking of vesicles to the distal appendages, components of Intraflagellar transport (IFT) are recruited in a TTBK2-dependent manner (Goetz *et al*, 2012; Tanos *et al*, 2013) to facilitate the growth of the ciliary axoneme by transporting various cargoes between the cilia base and tip (Prevo *et al*, 2017). Noteworthy, no evidence for a role in cilia formation has yet been provided for TTBK1, with its kinase domain highly similar to that of TTBK2 (Ikezu & Ikezu, 2014).

The Hedgehog pathway is perhaps the best-characterized signaling pathway relying on primary cilia. In the absence of HH ligand, the Patched (PTCH) receptor localizes inside the primary cilium and prevents ciliary accumulation of the receptor Smoothened (SMO). The binding of the HH ligand to PTCH initiates the removal of PTCH from the cilium, in turn leading to the accumulation of SMO inside cilia. SMO then interacts with cholesterol inside the cilium to switch the processing of GLI from its repressive form (GLI-R) to the active (GLI-A) form. Both GLI-R and GLI-A are translocated from primary cilia to the cell nucleus to repress and induce HH target genes, respectively (Anvarian *et al*, 2019; Radhakrishnan *et al*, 2020; Bangs & Anderson, 2017). The transcripts regulated by HH include components of the pathway (i.e. *PTCH1, GLI1*), as well as transcription factors regulating proliferation and cell fate decisions (Ingham, 2022).

The HH signaling pathway plays a prominent role in neural development. Its activity is critical for the establishment of the floor plate (the source of HH ligand in the ventral part of the neural tube) and the specification of individual neuronal types (Chiang *et al*, 1996; Briscoe *et al*, 1999). This activity is opposed by WNT/beta-catenin and BMP pathways. Thus, the neural tube pattern formation results from counteracting activities of HH, WNT, and BMP, regulated both spatially and temporally (Jessell, 2000; Ulloa & Briscoe, 2007). In addition, HH signaling drives cell proliferation during neurodevelopment in mice (Blaess *et al*, 2006) and can promote cell division in both neural and non-neural cell types (Liu *et al*,1998; Rowitch *et al*, 1999).

A major obstacle to the understanding of the role of the primary cilium and cilium-associated pathways has been the absence of a suitable cellular system to model human phenotypes and mechanisms. Human pluripotent stem cells (hPSCs), encompassing embryonic stem cells (ESCs) and induced pluripotent stem cells (iPSCs), can self-renew and differentiate into all cell types of the human body (Thomson *et al*, 1998; Takahashi *et al*, 2007). Consequently, they hold great promise for modeling both physiological and pathophysiological aspects of human embryogenesis “in a dish” (Park *et al*, 2008; Shahbazi *et al*, 2016). Interestingly, undifferentiated hPSCs can assemble primary cilia and express HH pathway components (Kiprilov *et al*, 2008; Banda *et al*, 2015; Wu *et al*, 2010). In mice, primary cilia start to appear in the epiblast, following embryo implantation (Bangs *et al*, 2015). However, the relevance of primary cilia for hPSCs self-renewal or differentiation capabilities is incompletely understood, with contradictions in the literature. While primary cilia have been proposed as instrumental for neural differentiation of hPSCs (Jang *et al*, 2016), their ablation, either in mouse embryos or in hPSCs, leads to surprisingly subtle neural differentiation defects (Tong *et al*, 2014; Cruz *et al*, 2022).

Events of neural differentiation and neural tube development can be modeled using hPSCs-derived neural rosettes, assemblies of radially organized neuroepithelial cells with a central lumen (Elkabetz *et al*, 2008; Fedorova *et al*, 2019). The neural rosettes, typically expressing early neuronal markers such as SOX2 and PAX6, can be readily specified into individual region-specific neuronal subtypes as well as serve as a progenitor niche (Kelava & Lancaster, 2016; Wilson & Stice, 2006; Elkabetz & Studer, 2008). HH pathway, together with Notch signaling, has been implicated in the maintenance of neural rosettes (Elkabetz *et al*, 2008).

Here, we combine the pharmacological activation of the HH signaling pathway with the CRISPR/Cas9-mediated ablation of TTBK2, to investigate the role of primary cilia in the proliferation and neural differentiation of hPSCs. We demonstrate that while TTBK2 and primary cilia are not required for the self-renewal of hPSCs, primary cilia and HH signaling regulate the size of hPSCs-derived neural rosettes. In addition, our data identify a role for TTBK1 in the cilium assembly pathway.

## Results

### HH signaling increases the size of neural rosettes

To examine the role of primary cilia-related HH signaling in hPSCs (human pluripotent stem cells), we treated CCTL14 hPSC line with Smoothened Agonist (SAG) (5nM, day (D)1-9) during neural rosettes differentiation (Fig. 1A). First. we observed ARL13B positive primary cilia pointing out of the apical cell surface (lumen) of the neural rosette (Fig. 1B), in agreement with a previous report (Banda *et al*, 2015). Next, we found that SAG treatment led to an induction of mRNA expression of HH target genes *GLI1* and *PTCH1* (Fig. 1C). In contrast, SAG did not have a notable effect on the expression of *NANOG, OCT4, GATA6, PAX6*, and *ISL1* mRNA (Suppl. Fig. 1A).

**Figure 1.**
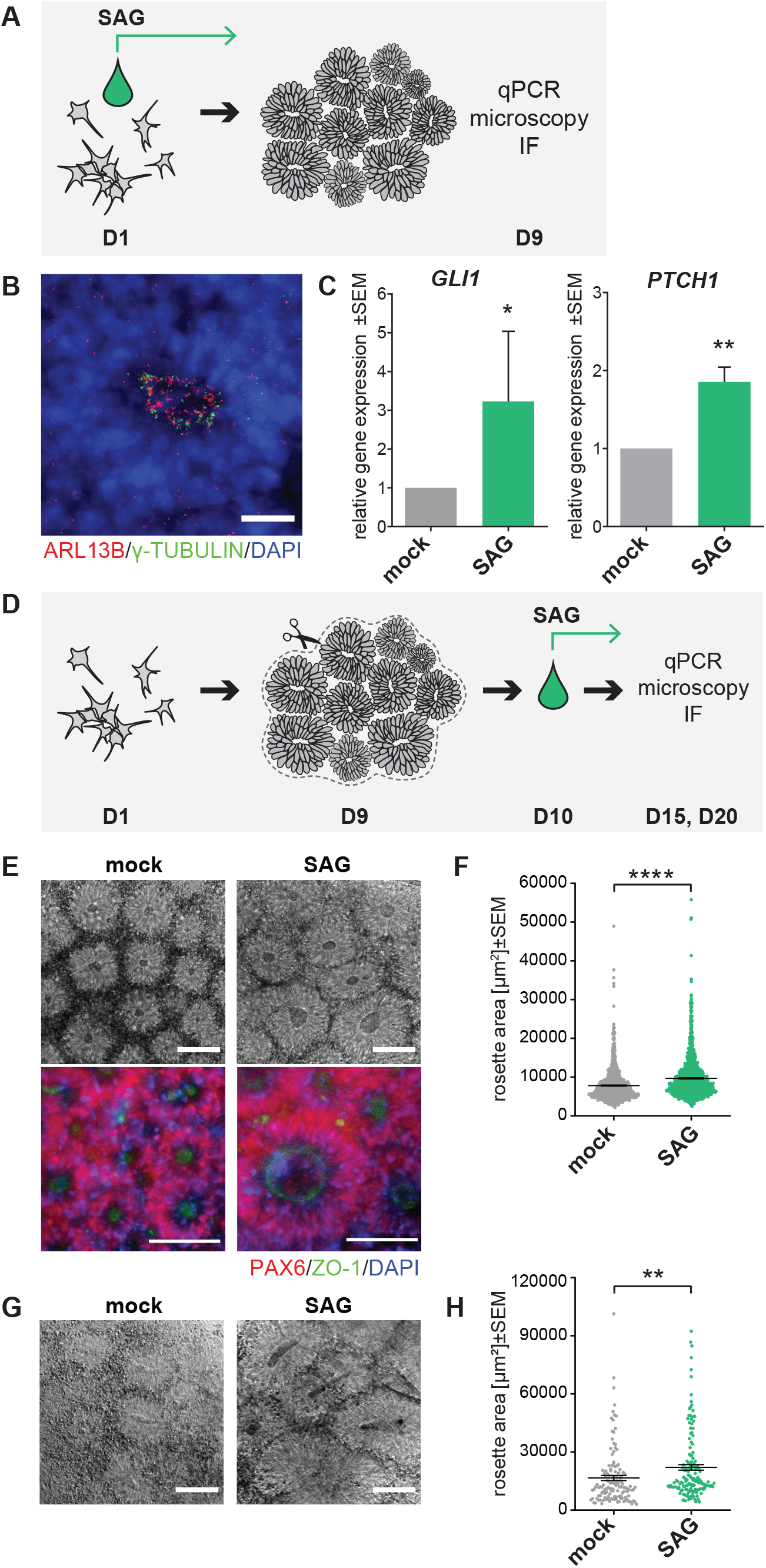
HH signaling increases the size of neural rosettes. A) Experimental design of neural rosettes early phase differentiation experiments. Cells were seeded on D0, SAG treatment started on D1 and cultures were analyzed on D9. B) Representative image of cilia IF staining in early phase differentiation of CCTL14 rosettes on D9, ARL13B staining was used to detect primary cilia, γ-TUBULIN staining indicates centrioles; scalebar=20μm. C) SHH target gene *GLI1* and *PTCH1* mRNA expression (qRT-PCR) in mock- and SAG-treated early phase of differentiation CCTL14 rosettes (D9); n=5, t-test. D) Experimental design of neural rosettes later phase differentiation experiments. D0-start of the experiment, D9 - patches of neural rosettes were transferred in a fresh culture plate, D10 – start of daily SAG treatment, D15/20 - analysis. E) Representative images of mock- or SAG-treated later stage differentiation CCTL14 rosettes on D15 in phase contrast (top) and IF (bottom), PAX6 staining was used to show ongoing neural differentiation, ZO1 staining indicates polarization in the lumen of neural rosettes; scalebar=100μm. F) Measurement of the rosette area in mock- or SAG-treated later stage differentiation CCTL14 rosettes on D15; n=3, t-test. G) Representative images of mock- or SAG-stimulated later stage differentiation Neo1 rosettes on D15 in phase contrast; scalebar=100μm. H) Measurement of the rosette area in mock- or SAG-treated later stage differentiation Neo1 rosettes on D15; n=4, t-test.

Next, we examined the effects of HH pathway activation in a later stage of neural rosette differentiation. To this end, we dissected patches of CCTL14-derived neural rosettes on D9 and cultured them in the presence of SAG/vehicle until D15 or D20 (Fig. 1D). On D15, cells forming the rosettes expressed neuronal maker PAX6 and showed tight junction protein ZO-1 highly enriched at the apical membrane (Fig. 1E), as expected (Hříbková *et al*, 2018). Consistent with the role of the HH pathway in neural tube patterning (Ulloa & Briscoe, 2007) we also observed increased expression of *SHH* mRNA and reduced expression of *WNT1* mRNA following the SAG treatment (Suppl. Fig 1B). Interestingly, we found the SAG-treated rosettes were notably larger than their mock-treated controls (Fig. 1E, F). Importantly, we confirmed the effect of SAG treatment on the neural rosette size using iPSC line Neo1 (Fig. 1G, H). In sum, our data confirm and extend the previous observation of a positive effect of HH pathway activation on neural rosette size (Elkabetz *et al*, 2008).

### TTBK2 is crucial for ciliogenesis but dispensable for self-renewal in hPSCs

Having validated our model system, we used CRISPR gene editing to establish TTBK2 knockout (KO) in CCTL14 hPSCs. We hypothesized that TTBK2 null hPSCs should be devoid of primary cilia. First, we verified that CRISPR-Cas9 successfully disrupted ORF of the TTBK2 locus in exon 4, which encodes a part of the kinase domain (Fig. 2A and Suppl. Fig. 2A). In total, we obtained 4 TTBK2 KO and 3 WT counterparts, which we used as controls in our following experiments. Of note, we also obtained one heterozygote line, with one allele disrupted and the other containing in-frame deletion within the kinase domain (Suppl. Fig. 2A), which we termed TTBK2 “mutant” (TTBK2 MUT). Next, we confirmed the lack of detectable levels of TTBK2 protein in total cell lysate (Suppl. Fig. 2B.) and at the mother centriole in TTBK2 KO/MUT lines (Fig. 2B and Suppl. Fig. 2C). Importantly, the lines lacking TTBK2 expression failed to form ARL13B+ primary cilia (Fig. 2B and Suppl. Fig. 2C).

**Figure 2.**
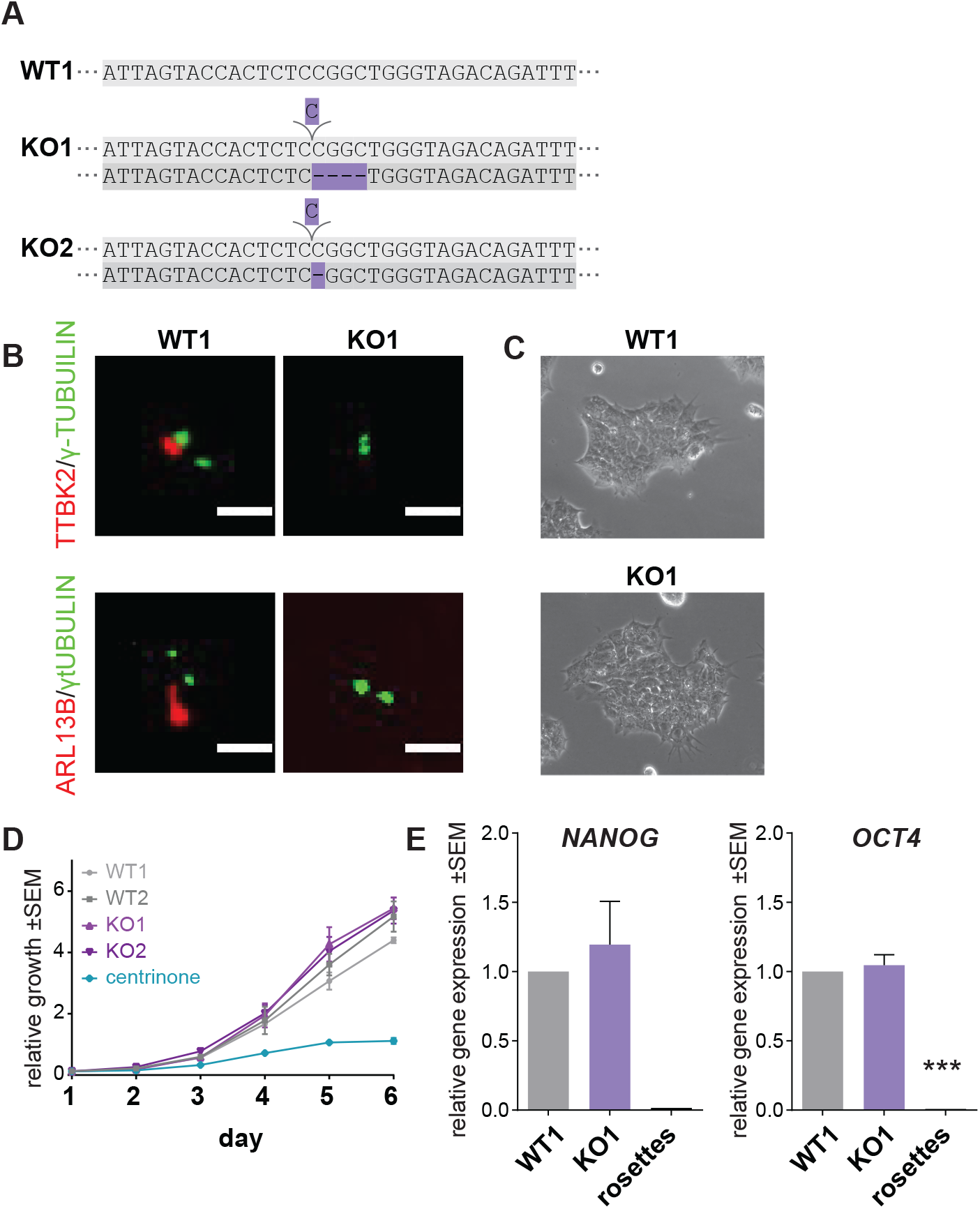
TTBK2 is crucial for ciliogenesis but dispensable for self-renewal in hPSCs. A) Schematic of the TTBK2 exon 4 sequence detail in representative WT and TTBK2 CRISPR cell lines, purple=insertion/deletion. B) Representative images of IF detection of TTBK2 (top) and cilia presence (bottom; visualized by ARL13B staining) in undifferentiated WT and TTBK2 KO, γ-TUBULIN staining was used to detect centrioles; scalebar=2μm C) Representative images of colony morphology of undifferentiated cells in WT1 and TTBK2 KO1. D) Relative growth comparison of indicated undifferentiated WT and TTBK2 KO lines assessed by crystal violet absorption measurement, centrinone treatment previously shown to impair the proliferation capacity was used as a control; n=3. E) mRNA expression (qRT-PCR) of pluripotency markers *NANOG* and *OCT4* in undifferentiated WT1 and TTBK2 KO1. Parental CCTL14 differentiated into neural rosettes was included for reference; n=4, one way ANOVA with Holm-Sidak’s multiple comparisons test.

Both TTBK2 WT and TTBK2 KO/MUT hPSCs grew in flat smooth-edged colonies with proto-typical hPSCs morphology (Fig. 2C, Suppl. Fig. 2D), which remained stable and self-renewing with no apparent signs of spontaneous differentiation in over 20+ passages. In line with that, we found that while depletion of centrioles by centrinone (150nM) treatment (Wong *et al*, 2015; Renzova *et al*, 2018) significantly impaired the proliferation capacity of hPSCs, the ablation of TTBK2/primary cilia showed no such effect (Fig. 2D and Supp. Fig. 2E). Moreover, TTBK2 WT and KO expressed comparable levels of pluripotency markers *OCT4* and *NANOG* (Fig. 2E, and Suppl. Fig. 2B and 2F). Similarly, we found no effect of SAG treatment on the proliferation of CCTL14 and Neo1 hPSCs (Suppl. Fig. 2G, H), the expression of pluripotency markers *OCT4* and *NANOG* (Suppl. Fig. 2I, J), and protein level of OCT4 (Suppl. Fig. 2K). These results confirm and extend recent observations on the primary cilia – self-renewal relationship (Cruz *et al*, 2022; Potjewyd *et al*, 2022) and suggest that TTBK2 and primary cilia are dispensable for self-renewal of hPSCs.

### Lack of TTBK2 increases the size of neural rosettes

Primary cilia are required for the correct activity of both the positive and negative HH pathway regulators (Ingham, 2022). As a result, primary cilia mutants show a loss of function HH phenotypes in some cell types and a gain of function HH phenotypes in others (Bangs & Anderson, 2017). Given that, we asked what specific function TTBK2/primary cilia serve in the regulation of the size of hPSC-derived neural rosettes. To this end, we adopted the protocol used earlier (Fig.1D). First, we confirmed that TTBK2 was absent from mother centrioles (Suppl. Fig. 3) and the primary cilia formation was impaired in TTBK2 KO neural rosettes (Fig. 3A, B). Next, we found both WT and KO TTBK2 lines were able to efficiently form neural rosettes expressing PAX6, with ZO-1 recruited to the apical membrane (Fig. 3C, D). Remarkably, we noticed that neural rosettes were notably larger in mock-treated TTBK2 KOs than in the corresponding WT controls. In addition, while SAG treatment promoted the rosette size increase in WT, it failed to show an additive effect in TTBK2 KO (Fig. 3D, E). Next, we asked if altered cell proliferation may explain the observed effects on neural rosette size. To this end, we examined the number of phospho histone 3 (pH3) positive cells in neural rosettes on D15. Indeed, we found the relative number of pH3+ cells was elevated in TTBK2 KO and MUT cells, respectively (Fig. 3F-G).

**Figure 3.**
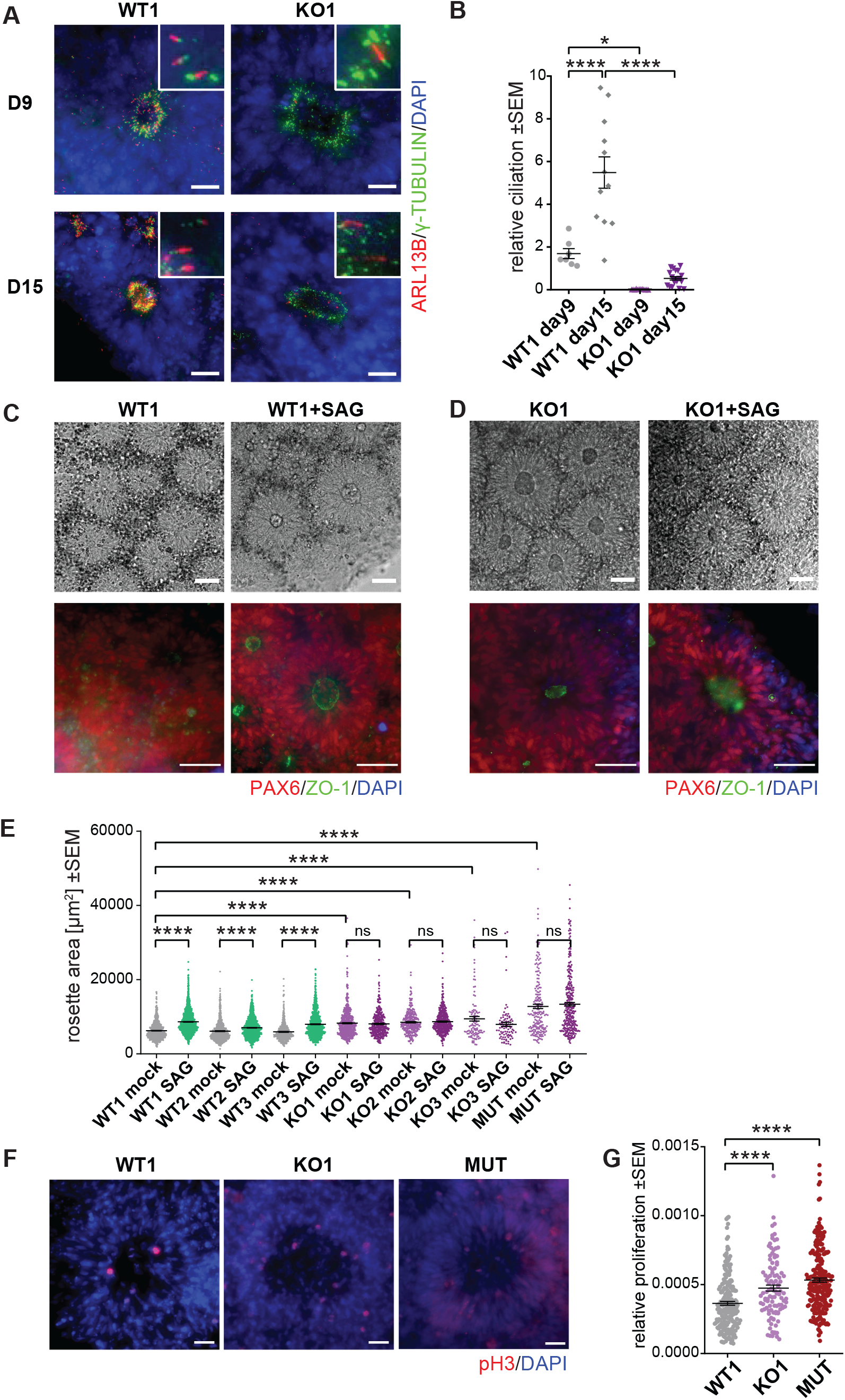
Lack of TTBK2 increases the size of neural rosettes. A) Representative images of IF detection of primary cilia (visualized by ARL13B staining) in WT1- and TTBK2 KO1-derived neural rosettes at D9 (top) and D15 (bottom), γ-TUBULIN was used to indicate centrioles; scalebar=20μm. B) Relative ciliation (measured as ARL13B positive rosette area fraction) in WT1- or TTBK2 KO1-derived neural rosettes at D9 and D15; n=2, one-way ANOVA with Holm-Sidak’s multiple comparisons test. C) Mock- or SAG-treated WT1-derived neural rosettes visualized in phase contrast (top) or IF (bottom). PAX6 staining was used to monitor ongoing neural differentiation, ZO1 staining visualizes polarization of the lumen; scalebar=50μm. D) Mock- or SAG-treated TTBK2 KO1-derived neural rosettes visualized in phase contrast (top) and IF (bottom), PAX6 staining was used to monitor ongoing neural differentiation, ZO1 staining visualizes polarization of the lumen; scalebar=50μm. E) Measurement of the rosette area in mock- or SAG-treated WT- and TTBK2 KO-derived neural rosettes, respectively, on D15; n=3, one-way ANOVA with Holm-Sidak’s multiple comparisons test. F) Representative images of Phospho-histon3 (pH3) IF detection in the indicated conditions on D15; scalebar=20μm. G) Relative proliferation as a ratio of pH3 positive cells number per rosette area on D15; n=3, one-way ANOVA with Holm-Sidak’s multiple comparisons test.

In sum, these results demonstrate that while TTBK2 and primary cilia are required for the response to HH pathway stimulation in neural rosettes, their ablation mimics the HH pathway activation phenotype.

### Tau tubulin kinase 1 (TTBK1) regulates cilia formation

Our data indicated that TTBK2 is indispensable for ciliogenesis in undifferentiated hPSCs (Fig. 2B, Suppl. Fig.2C), in agreement with its role in primary cilia formation in other biological systems (Goetz *et al*, 2012; Bernatik *et al*, 2020). Surprisingly, our data also revealed that even when TTBK2 was absent (Suppl. Fig. 3), some primary cilia still formed with the ongoing differentiation of TTBK2 KO-derived neural rosettes (Fig. 3A), albeit at a much-reduced rate over WT (Fig. 3B). We hypothesized that the absence of TTBK2 may be partially rescued specifically during neural differentiation. In turn, we considered Tau tubulin kinase 1 (TTBK1) as a plausible mediator of such rescue effect. As we already mentioned, TTBK1 had no previous links to the regulation of ciliogenesis, but shared a high degree of sequence homology in its kinase domain with TTBK2 (Suppl. Fig. 4C), and showed high expression levels in the CNS (Ikezu & Ikezu, 2014).

First, we examined the expression of *TTBK1* and found its mRNA levels significantly upregulated in differentiating CCTL14 hPSC - derived neural rosettes (Fig. 4A), and in neuro-differentiated cultures and organoids of i3N iPSCs (Suppl. Fig. 4A). Next, we used TTBK1/2 inhibitor (Potjewyd *et al*, 2022) (1μM) during neural rosette differentiation (D12-15) of TTBK2 WT and KO hPSC, respectively. We found that the treatment not only reduced ciliogenesis in TTBK2 WT but almost fully eliminated cilia formation in TTBK2 KO (Fig. 4C). Importantly, the inhibition of TTBK1/2 led to an additional increase of the rosette size in TTBK2 KO (Fig. 4D). To corroborate this observation, we transiently expressed TTBK1-Halo in TTBK2 KO hPSC and hTERT-RPE-1 TTBK2 KO cells. Remarkably, we found that while transiently expressed GFP-TTBK2 localized to basal bodies, TTBK1-Halo failed to do so, nevertheless its expression showed a rescue effect on the formation of ARL13b+ primary cilia (Fig. 4E, F, Suppl. Fig. 4B). In sum, these data suggest that TTBK1 participates in the regulation of primary cilia assembly and that ciliogenesis in hPSC-derived neural rosettes is under the control of both TTBK1 and TTBK2.

**Figure 4.**
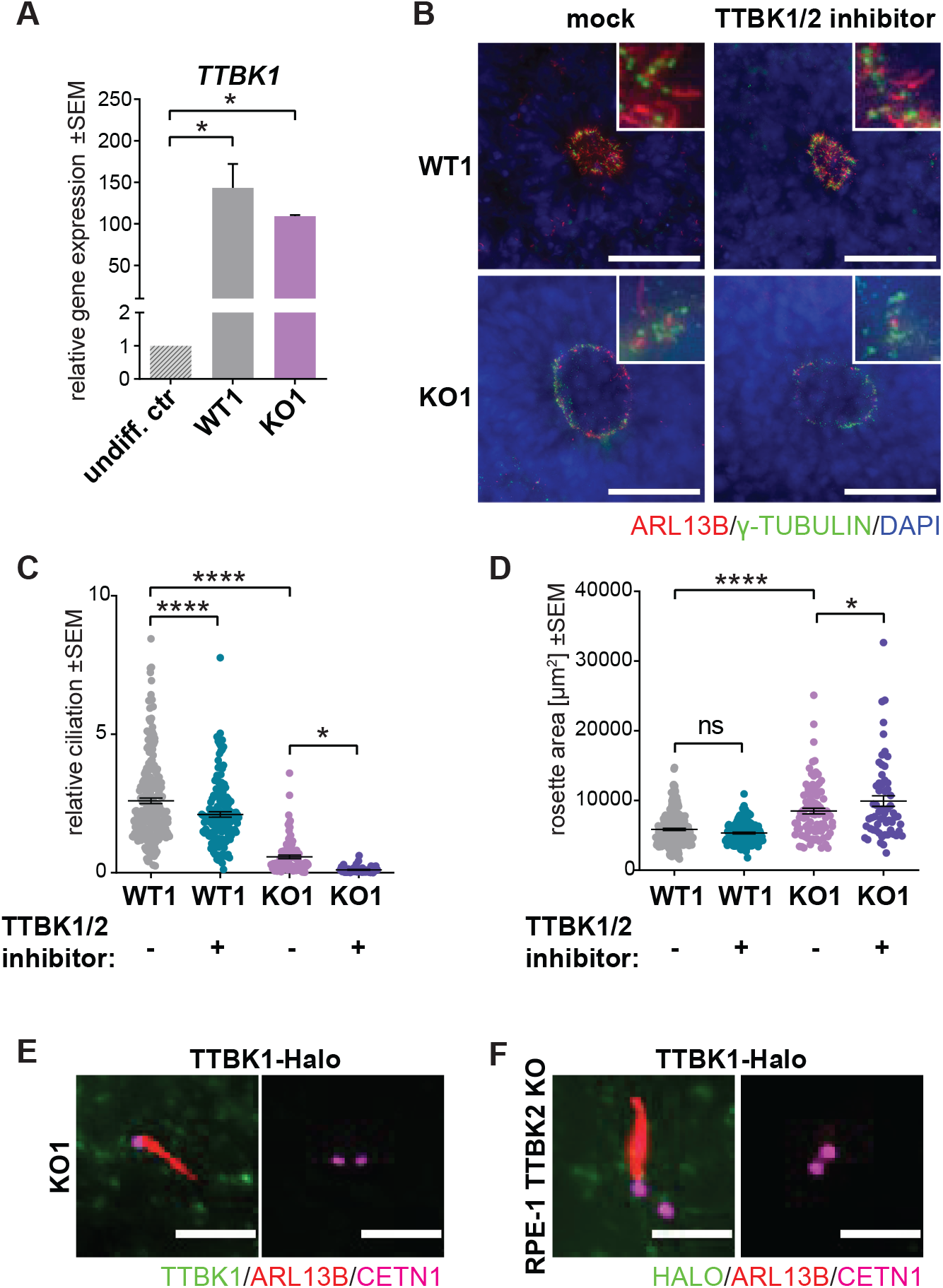
Tau tubulin kinase 1 (TTBK1) regulates cilia formation. A) mRNA expression (qRT-PCR) of *TTBK1* in undifferentiated parental cell line CCTL14, and WT1- or TTBK2 KO1-derived neural rosettes on D15; n=2-3, one-way ANOVA with Holm-Sidak’s multiple comparisons test. B) IF detection of primary cilia (visualized by ARL13B staining) in mock- or TTBK1/2 inhibitor-treated WT1 and TTBK2 KO1, respectively, on D15, γ-TUBULIN was used to detect cetrioles; scalebar=50μm. C) Relative ciliation (measured as ARL13B positive rosette area fraction) in WT1- or TTBK2 KO1-derived neural rosettes, treated as indicated, on D15; n=2, one-way ANOVA with Holm-Sidak’s multiple comparisons test. D) Rosette area measurement in WT1- or TTBK2 KO1-derived neural rosettes, treated as indicated, on D15; n=2, one-way ANOVA with Holm-Sidak’s multiple comparisons test. E) IF detection of primary cilia (visualized by ARL13B staining) in TTBK2 KO1 transfected with TTBK1-Halo (left) or not transfected (right). CETN1 staining was used to detect centrioles; scalebar=3μm. F) IF detection of primary cilia (visualized by ARL13B staining) in hTERT RPE-1 TTBK2 KO cells transfected with TTBK1-Halo (left) or not transfected (right), CETN1 was used to indicate centrioles; scalebar=3μm.

## Discussion

Both primary cilia and stem cells have emerged as key regulators of embryo development and tissue homeostasis. Here we have explored the functions of primary cilia using a panel of WT and CRISPR/Cas9-edited hPSCs devoid of TTBK2 and, in turn, primary cilia. While we found no major function for TTBK2/primary cilia in hPSCs self-renewal, we identified a role for primary cilia and HH pathway in the regulation of hPSCs –derived neural rosettes. In addition, our data implicated TTBK1 in the regulation of ciliogenesis in human cells.

Primary cilia, capable to transduce HH signal, were reported in hPSCs previously, but with unclear biological significance (Kiprilov *et al*, 2008; Banda *et al*, 2015). Our data indicate that TTBK2 and primary cilia do not play a major role in the regulation of undifferentiated hPSCs. In support of our observation, activation of the HH pathway failed to prevent hPSCs spontaneous differentiation following FGF2-withdrawal (Wu *et al*, 2010). While we cannot completely rule out the impact of specific culture conditions, we conclude that hPSCs lacking primary cilia can be efficiently propagated without notable changes in the morphology of individual colonies or pluripotency markers expression. When finalizing this manuscript, similar results have been reported using KIF3A/KIF3B knockout hPSCs (Cruz *et al*, 2022) or TTBK2 knockout iPSCs (Potjewyd *et al*, 2022).

So why would hPSCs invest their resources to assemble the primary cilium, if this organelle is not crucial at this particular cell stage? We speculate the presence of primary cilia in the undifferentiated cell state may facilitate the later execution of a specific differentiation program (Anderson & Stearns, 2009). In a way, this may be conceptually similar to the poised state of many gene promoters in hPSCs, thus subsequently allowing an efficient response upon appropriate stimuli (Macrae *et al*, 2022).

Our data indicate that ablation of primary cilia in hPSCs leads to a similar phenotype as the activation of the HH pathway – increased size of neural rosette. This observation is in agreement with a different dependency of individual HH activators and inhibitors on the presence of primary cilia. Interestingly, while various HH “gain of function” phenotypes have been reported following the disruption of primary cilia (Bangs & Anderson, 2017), loss of primary cilia due to TTBK2 ablation in the neural tube in mice has been associated with HH “loss of function” defects (Goetz *et al*, 2012). In addition, disruption of ciliogenesis in OFD1 KO mouse embryonic stem cells led to reduced activation of the HH pathway in the course of neural differentiation (Hunkapiller *et al*, 2011). It remains to be determined what factors underlay such differences between mouse and human cells. Interestingly, recent work has postulated that hPSCs-derived neural rosettes are formed by a mechanism of secondary neurulation, which is in mice restricted to the most caudal parts of the developing neural tube (Fedorova *et al*, 2019).

Primary cilia were proposed as essential for hPSCs conversion into PAX6+ neural progenitors, hence for the neural fate acquisition (Jang *et al*, 2016). Our data challenge such a model and suggest that TTBK2 and primary cilia are not critical for the acquisition of PAX6+ neural progenitor fate during hPSC differentiation. Instead, primary cilia seem to regulate the proliferation of neural progenitors at the neural rosette stage. This observation is in agreement with a reported accumulation of SOX2+ neural progenitors in KIF3A and KIF3B KO hPSCs (Cruz *et al*, 2022) and with the absence of early neurodifferentiation phenotypes in mice with primary cilia defects (Huangfu *et al*, 2003; Goetz *et al*, 2012; Tong *et al*, 2014). Moreover, given the recent progress in the development of inhibitors of TTBK1/2 (Halkina *et al*, 2021; Potjewyd *et al*, 2022), temporal ablation of primary cilia could be exploited to tweak the yield of neurodifferentiation protocols.

TTBK1 and TTBK2 share almost 60% identity and 70% similarity in their kinase domains, making them the closest relatives within the CK1 kinase family (Ikezu & Ikezu, 2014). Previous data, including our own, established that TTBK2 is essential for primary cilia formation in several systems – no other kinase seemed able to compensate for a complete loss of TTBK2 (Goetz *et al*, 2012; Bernatik *et al*, 2020). Similarly, our current data show that the ablation of TTBK2 in undifferentiated hPSCs leads to a complete loss of primary cilia. Intriguingly, however, our results further suggest that TTBK1 can partially compensate for the absence of TTBK2 in hPSCs-derived neural rosettes. This is quite surprising, as the Proline-rich motif implicated in TTBK2-CEP164 interaction (Oda *et al*, 2014; Rosa e Silva *et al*, 2022; Čajánek & Nigg, 2014) and, in turn, the recruitment of the kinase to the mother centriole, is poorly conserved in TTBK1 (Suppl. Fig. 4D). One plausible explanation is that concentrating the kinase activity at mother centriole is not strictly necessary for the efficient phosphorylation of its key substrates, provided the levels and activity of the kinase outside of the mother centriole are sufficiently high. Indeed, undifferentiated hPSCs have low levels of TTBK1, not sufficient for any noticeable contribution to primary cilia assembly. In contrast, TTBK1 expression is significantly upregulated during neural rosette formation, and, consequently, TTBK1 becomes competent to affect ciliogenesis. Importantly, our rescue experiment with transient TTBK1 transfection represents proof of concept that TTBK1 is able to regulate primary cilia. Our model is attractive also from the point of TTBK2 frame-shift mutations, associated with spinocerebellar ataxia 11 (Houlden *et al*, 2007). Here, the resulting truncated protein moieties lack the C-terminal CEP164 binding motif and hence are considered unable to support ciliogenesis (Goetz *et al*, 2012; Bowie *et al*, 2018). In addition, primary cilia are emerging as critical regulators of CNS functionality (Bowie & Goetz, 2020; Schmidt *et al*, 2022; Sheu *et al*, 2022). Given that TTBK1 has been considered a plausible therapeutic target for the treatment of Alzheimer’s disease (AD) (Ikezu *et al*, 2020; Halkina *et al*, 2021; Nozal & Martinez, 2019; Taylor *et al*, 2018), its direct involvement in primary cilia regulation may significantly hamper these efforts in AD targeting. Therefore, future studies should address in which cell types is TTBK1 able to regulate ciliogenesis and the exact mechanism of its action.

## Material and Methods

### Cell culture and transfection

CCTL14 cells https://hpscreg.eu/cell-line/MUNIe007-A were cultured in the undifferentiated state as a feeder-free monolayer on Matrigel (Corning)-coated plastic in daily changed MEF-conditioned hESC medium ((DMEM/F12, 15% Knockout serum replacement (both from Gibco), 2 mM L-glutamine, 1x nonessential amino acids (NEAA) (both from Biosera), ½x Zell Shield (Minerva Biolabs), 100 μmol/L β-mercaptoethanol (Sigma-Aldrich) and 10 ng/ml hFGF2 (Invitrogen)). Conditioned medium was enriched with 2mM L-glutamine, ½x Zell Shield, and 10 ng/ml hFGF2 before use. iPSC Neo1 w1 cells (Renzova *et al*, 2018) were cultured in the undifferentiated state as a feeder-free monolayer on Matrigel (Corning)-coated plastic in daily changed complete mTeSR medium (StemCell Technologies) with ½x Zell Shield. Cells were regularly passaged using Tryple Express (Thermo Fisher Scientific). Where indicated, undifferentiated cells were treated with 500 nM SAG (Sigma) on D1 and D2 after seeding and analyzed on D3. i3N cells were maintained and differentiated as described in detail in (Fernandopulle *et al*, 2018). hTERT RPE-1 KOs were cultured in DMEM/F12, 10% FBS, and 2 mM L-glutamine as described in (Bernatik *et al*, 2020).

For transient transfection experiments, TTBK2 KO cell lines were seeded on glass coverslips (Matrigel-coated for hPSCs) and the next day transfected with 0.5μg TTBK1-HaloTag^®^ human ORF in pFN21A (FHC12512, Promega) or pglap1-TTBK2 (“GFP-TTBK2”, (Bernatik *et al*, 2020)) using Lipofectamine 3000 (Invitrogen). The medium was changed 4h after transfection for fresh culture medium. To promote ciliogenesis, hTERT RPE-1 24h were starved in serum-free DMEM/F12 with 2 mM L-glutamine 24h after transfection for 24h.

### Neural rosette differentiation

Cells were seeded in density 4000-6000 cells/cm^2^ in a complete growth medium with 20μM ROCK1 inhibitor Y-27632 (Selleckchem). On the next day (D1) medium was changed for Rosette differentiation medium (DMEM/F12:Neurobasal 1:1 (both from Gibco), N2, B27 with vitamin A (both from ThermoFisher), 2 mM L-Glutamine, 1x NEAA, ½x Zell Shield) with 20μM Y-27632, and 20μM TGF-β inhibitor SB 431542 (Sigma). Where indicated, 5nM SAG (Sigma) treatment started on D1. On D4 the medium was changed for Rosette differentiation medium (without inhibitors), the medium was changed daily until D9. Around D6 rosettes started to appear. For the early stage of the differentiation experiment (Fig. 1A), the cells were fixed/harvested on D9. For later stages of neural rosette differentiation (Fig. 1D), clusters of rosettes were cut out on D9 and transferred on a Matrigel-coated plate (for immunostaining glass coverslips were inserted in the wells and coated with Matrigel) with fresh Rosette differentiation medium. On D10, the medium was changed, and 5nM SAG was added, where indicated. Half of the medium (with vehicle/SAG) volume was replaced daily until D15. Where indicated, rosette cultures were treated with TTBK1/2 inhibitor (1μM) (gift from Alison Axtman (Potjewyd *et al*, 2022)) daily on D12-D15.

### Phase contrast imaging

Phase contrast images of differentiated rosettes were acquired on D15 using inverted microscope LEICA DM IL LED equipped with Leica DFC295 camera (Leica Microsystems), Leica Application Suite (LAS) Version 3.8.0, and HI PLAN I 10x/0.22 objective. Freehand selection tool in Fiji (version 2.1.0, (Schindelin *et al*, 2012) was employed to outline the rosette outer border, and the area inside the selected outline was measured as the Rosette area.

### Immunofluorescence (IF) imaging

For cilia staining, the cultures were fixed by −20°C methanol, washed in PBS, blocked in blocking buffer (1% BSA in PBS), and stained 1h/RT or overnight/4°C with the following primary antibodies in blocking buffer: ARL-13B (rabbit, 17711-1-AP, Proteintech, 1:1000), ARL13B (mouse, sc-515784, Santa-Cruz, 1:200), γ-tubulin (mouse, T6557, Sigma, 1:1000), TTBK1 (mouse, HPA031736, Sigma, 1:500), TTBK2 (rabbit, HPA018113, Sigma, 1:500), Halo-tag (mouse, 28a8 ChromoTek, 1:200), Alexa Fluor 647-direct labeled CETN1 (rabbit, 12794-1-AP, Proteintech/ A20186, Invitrogen, 1:50). For neural rosette markers and phosphoH3 staining, the cultures were fixed in 4% PFA 20min/4°C, washed 2x in PBS, permeabilized in 0,2% Triton-X100 in PBS 5min/RT, and stained 1h/RT or overnight/4°C with following primary antibodies in 0,2% Triton-X100 in PBS: PAX6 (rabbit, CS60433, Cell Signaling, 1:300), ZO-1-1A12 (mouse, 33-9100, Invitrogen, 1:800), Phospho-Histone H3 Thr11 (rabbit, #9764 Cell Signaling, 1:300). Coverslips were then washed 3x 5min in PBS and stained with donkey-raised secondary antibodies in either blocking buffer or 0,2% Triton-X100 in PBS, 1h/RT: anti-mouse or anti-rabbit Alexa Fluor 488, 568 or 647 (all Invitrogen, 1:1000). Nuclei were stained with DAPI or Hoechst 5min/RT. After washing, coverslips were mounted using Glycergel (Agilent) or ProLong™ Glass Antifade Mountant (Invitrogene).

Z-stack images were acquired on Zeiss AxioImager.Z2, with Hamamatsu ORCA Flash 4.0 camera, using 40x/1.3 Plan Apochromat OIL or 63x/1.4 Plan-Apochromat OIL objective controlled by Zen Blue Software.

### Image analysis

Z-slices were projected in one layer by Maximal Intensity Z-projection, a Color composite image was created in Fiji. To approximate the amount of the cilia (high density of the cilia in rosette lumen hampers counting their exact numbers), we measured the ARL13B positive rosette area fraction. The freehand selection tool in Fiji was employed to outline the rosettes outer borders, which were then added to the ROI manager. The image was split into separate color channels and in the channel where the ARL13B signal was acquired a threshold mask was applied and adjusted manually to cover the area with the ARL13B-stained cilia. The fraction of the rosette (%) covered by the mask was then measured as an ARL13B-positive rosette area fraction. To quantify the proliferation of the cells in rosettes, we counted phospho-histon3 (pH3)-positive cells per rosette. The freehand selection tool in Fiji was again used to outline the rosette outer borders. The pH3-positive cells within the selected outline were manually counted and a ratio of the pH3 cell number to the rosette area was obtained.

### qRT-PCR

Total RNA was isolated using RNA blue reagent according to the manufacturer’s recommendations. 0.3-1ug RNA was used for cDNA synthesis using Transcriptor First Strand cDNA Synthesis Kit (Roche). 2ul of 8x diluted cDNA was used in 10ul qPCR reactions with LightCycler^®^ 480 SYBR Green I Master with primers listed in Table 1. according to the manufacturer’s protocol and monitored in real-time using LightCycler^®^ 480 Instrument II (Roche). Relative gene expression was calculated using 2^-ΔΔCq^ method; GAPDH was used as the reference gene.

**TABLE 1:**
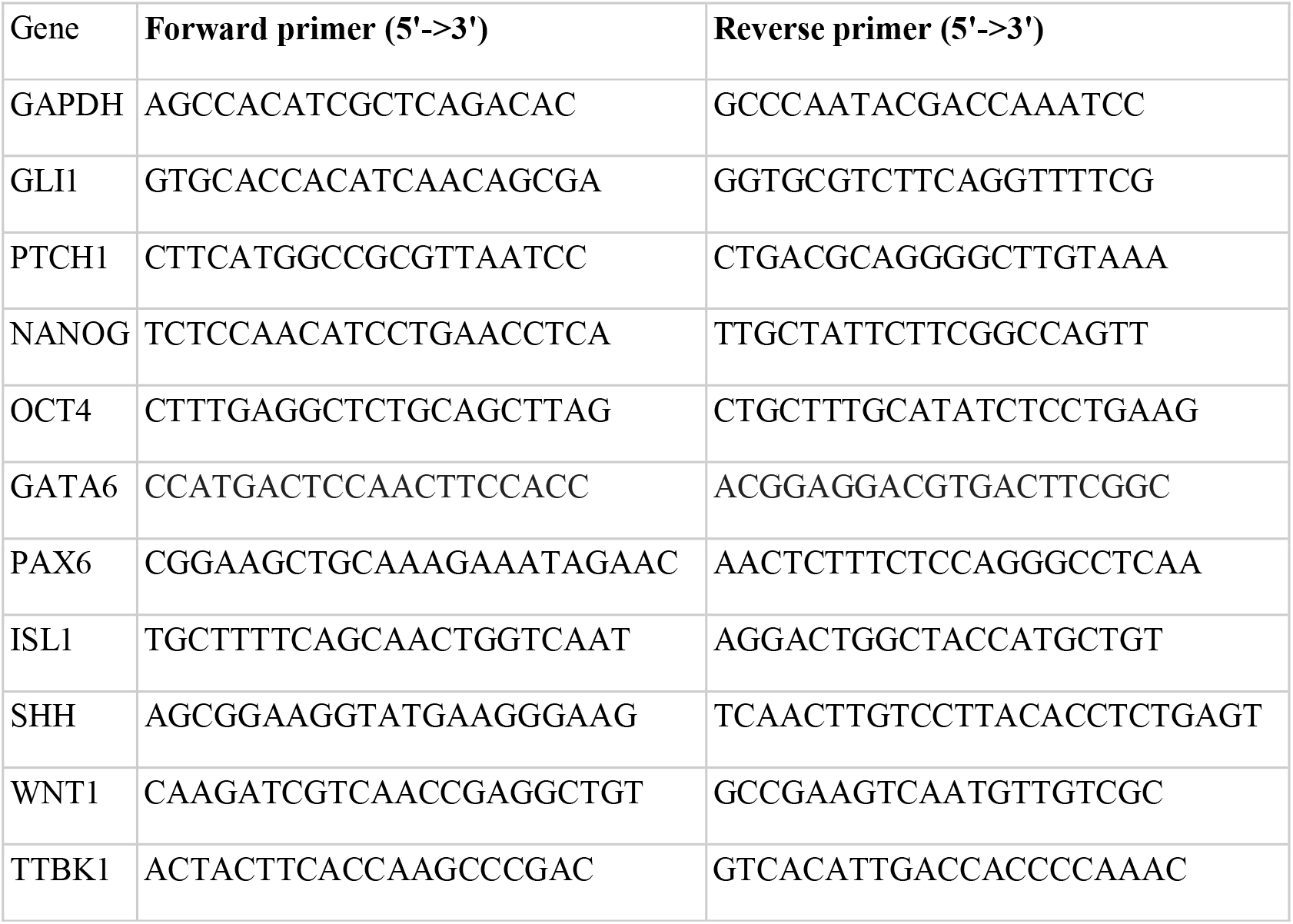
qRT-PCR Primers sequences

### Western Blot

Protein expression was analyzed by western blotting as described before (Renzova *et al*, 2018). Briefly, cells were lysed in SDS lysis buffer (50 mM Tris-HCl pH 6.8, 10% glycerol, 1% SDS). Protein concentration was measured using DC Protein Assay Kit (Bio-Rad) and an equal amount of protein was loaded in 8% gel. The following primary antibodies were used: TTBK2 (rabbit, HPA018113, Sigma, 1:250), NANOG (rabbit, 3580S, Cell Signaling, 1:1000), OCT4 (mouse, SC-5279, Santa Cruz, 1:4000), α-TUBULIN (mouse, 6631-1 Proteintech, 1:2000) was used as a loading control. To detect the signal, anti-rabbit (7074s, Cell Signaling, 1:3000), or anti-mouse (A4461, Sigma, 1:3000) secondary HRP-conjugated antibodies were used.

### Growth curves

The relative growth rate was determined using a crystal violet assay as described before (Renzova *et al*, 2018). Briefly, cells were seeded on 96 well plates, and mock- or SAG (500nM) treated from D1. Cultures were fixed on D1-6 in 4% formaldehyde, PBS washed, and incubated in 0.5% crystal violet (1 hour). Following 3x H2O wash, they were incubated in 33% acetic acid (20 minutes/shaking). Relative growth was determined as an increase in absorbance at 570 nm.

### CRISPR/Cas9 knock-out

TTBK2 gRNA (Bernatik *et al*, 2020) cloned into pSpCas9 (BB)-2A-GFP vector was repeatedly transfected into CCTL14 line, as described before (Bohaciakova *et al*, 2017). pSpCas9(BB)-2A-GFP vector was a gift from Feng Zhang (https://www.addgene.org/48138, (Ran *et al*, 2013)). GFP-positive cells were single-cell sorted using BD FACS ARIA II to matrigel-coated 96-well plates. Selected clones were verified for TTBK2 presence by immunostaining, expanded, and sequenced to verify the disruption of the TTBK2 locus.

### Sequence alignment

Jalview software (version 2.11.2.5) (Waterhouse *et al*, 2009) was used for TTBK1 and TTBK2 sequence alignment visualization.

### Statistical analysis

Quantitative data are presented as the mean□±□SEM. Statistical differences among groups were evaluated by t-test or one-way analysis of variance (ANOVA) followed by Holm-Sidak’s or Tukey’s multiple comparisons tests. For all statistical analyses, P value <0.05 was considered significant (*P□<□0.05, **P□<□0.01, ***P□<□0.001, and ****P□<□0.0001). All statistical analyses were performed with GraphPad Prism (GraphPad Software; www.graphpad.com). All experiments were performed at least in triplicate unless stated otherwise.

## Acknowledgment

We thank Alison Axtman, Jan Raška, and Tomáš Bárta for sharing reagents or samples, Tomáš Loja for help with FACS, and Hana Hříbková with Dáša Bohačiaková for advice on neural rosette differentiation protocol. The work was supported by grants from the Czech Science Foundation (22-13277S) and the Faculty of Medicine (MUNI/11/SUP/03/2022) to L.C. We acknowledge the core facility CELLIM supported by the Czech-BioImaging large RI project (LM2018129 funded by MEYS CR) for their support with obtaining scientific data presented in this article.

## Author contributions

L.B. – performed experiments and analyzed data, L.C. – conceived and supervised the project, and together with L.B. wrote the paper.

## Figure legends

**Supplementary Figure 1. SAG treatment in neural rosette differentiation does not affect selected pluripotency and differentiation markers**

A) mRNA expression (qRT-PCR) of selected markers in mock- and SAG-treated CCTL14 rosettes on D9; n=5, one-way ANOVA with Tukey’s multiple comparisons test.

B) mRNA expression (qRT-PCR) of selected markers in mock- and SAG-treated CCTL14 rosettes on D20; n=2, t-test.

**Supplementary Figure 2. TTBK2 is crucial for ciliogenesis but dispensable for selfrenewal in hPSCs and SAG treatment does not affect pluripotency in undifferentiated hPSC**

A) Schematic of the TTBK2 exon 4 sequence detail in WT and TTBK2 KO/MUT cell lines, purple=insertion/deletion, dark red=in frame deletion (mutant).

B) Representative images of western blot detection of TTBK2, NANOG and OCT4 protein expression in undifferentiated WT and TTBK2 KO/MUT lines; α-TUBULIN was used as a loading control.

C) Representative images of IF detection of TTBK2 (top) and primary cilia (bottom; visualized by ARL13B staining) in undifferentiated WT and TTBK2 KO/MUT lines, γ-TUBULIN staining was used to detect centrioles; scalebar=2μm.

D) Representative image of colony morphology of undifferentiated cells in TTBK2 MUT line.

E) Relative growth comparison of indicated undifferentiated WT and TTBK2 KO/MUT lines assessed by crystal violet absorption measurement, centrinone treatment previously shown to impair the proliferation capacity was used as a control; n=3.

F) mRNA expression (qRT-PCR) of pluripotency markers *NANOG* and *OCT4* in indicated undifferentiated WT and TTBK2 KO/MUT lines; n=4, one-way ANOVA with Holm-Sidak’s multiple comparisons test.

G) Relative growth comparison of mock- and SAG-treated WT CCTL14 line assessed by crystal violet absorption measurement, centrinone treatment previously shown to impair the proliferation capacity was used as a control; n=3.

H) Relative growth comparison of mock- and SAG-treated WT Neo1 line assessed by crystal violet absorption measurement, centrinone treatment previously shown to impair the proliferation capacity was used as a control; n=3.

I) mRNA expression (qRT-PCR) of pluripotency markers *NANOG* and *OCT4* in mock- and SAG-treated (48h) WT undifferentiated CCTL14 line, CCTL14 differentiated into neural rosettes was included for reference; n=3, one-way ANOVA with Tukey’s multiple comparisons test.

J) mRNA expression (qRT-PCR) of pluripotency markers *NANOG* and *OCT4* in mock- and SAG-treated (48h) WT undifferentiated Neo1 line, CCTL14 differentiated into neural rosettes was included for reference; n=3, one-way ANOVA with Tukey’s multiple comparisons test.

K) Representative images of western blot detection of pluripotency marker OCT4 in mock- and SAG-treated (48h) WT undifferentiated CCTL14 and Neo1; α-TUBULIN was used as a loading control.

**Supplementary Figure 3. TTBK2 is not expressed in TTBK2 KO-derived neural rosettes**

Representative images of IF detection of TTBK2 presence in WT1- and TTBK2 KO1-derived neural rosettes on D9 (top) and D15 (bottom), γ-TUBULIN was used to detect centrioles; scalebar=20μm.

**Supplementary Figure 4. Tau tubulin kinase 1 (TTBK1) regulates cilia formation**

A) mRNA expression (qRT-PCR) of *TTBK1* in undifferentiated parental cell line i3N compared to differentiated neurons on D17 and organoid on D85; n=2, one-way ANOVA with Holm-Sidak’s multiple comparisons test.

B) Representative images of IF detection of primary cilia (visualized by ARL13B staining) in TTBK2 KO1 cells transfected with GFP-TTBK2 (left) or not transfected (right), γ-TUBULIN was used to detect centrioles, scalebar=3μm.

C) N-terminal TTBK1 and TTBK2 kinase domains alignment, identical amino acids are shown in dark blue.

D) C-terminal CEP164-binding region in TTBK2 (amino acids 842-1244) aligned to TTBK1, identical amino acids are shown in dark blue, Proline-rich motif necessary for CEP164 binding is highlighted in magenta.

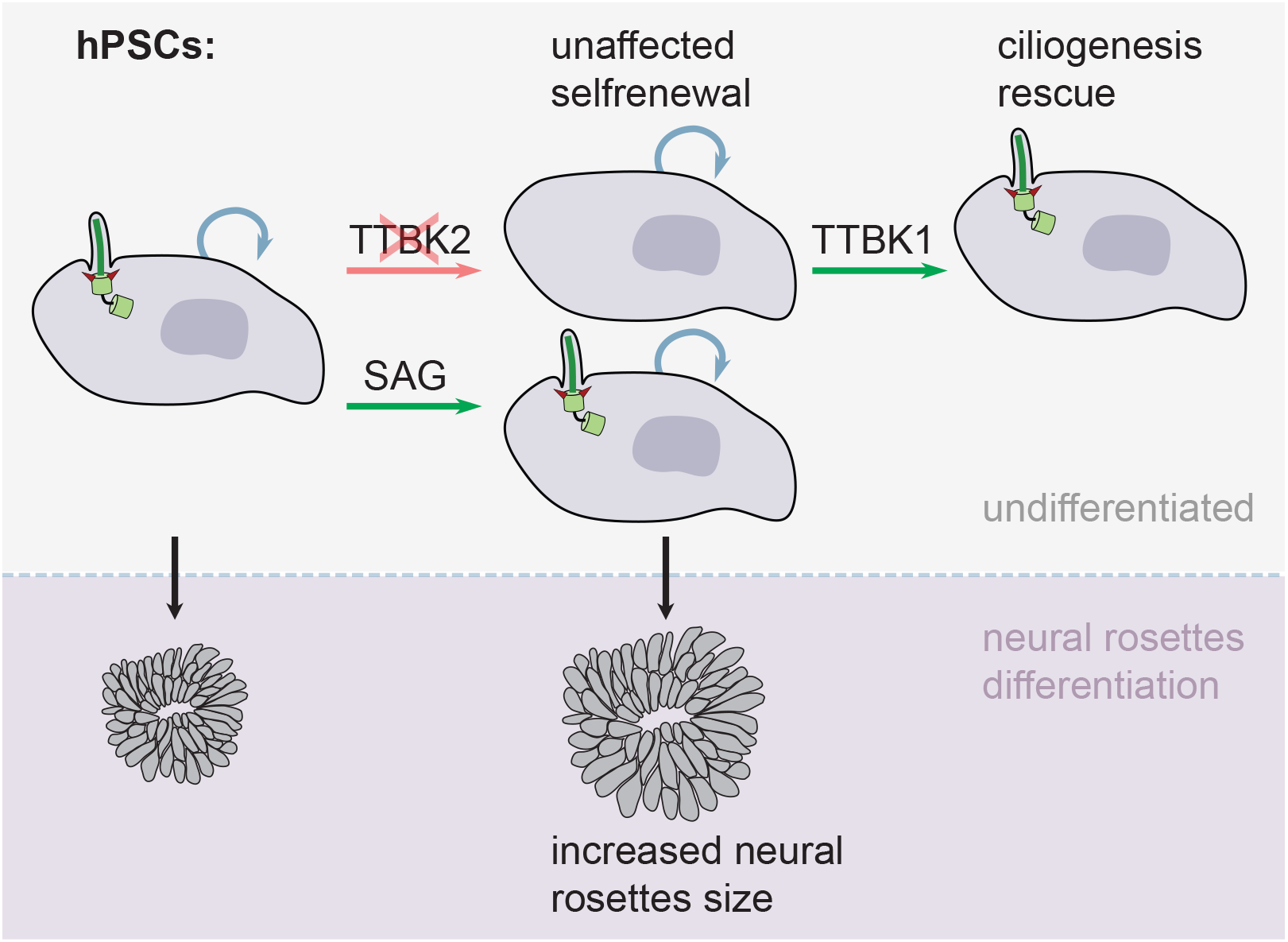

