## Supplementary Figure 1 for "Tau tubulin kinase 1 and 2 regulate ciliogenesis and human pluripotent stem cells–derived neural rosettes"

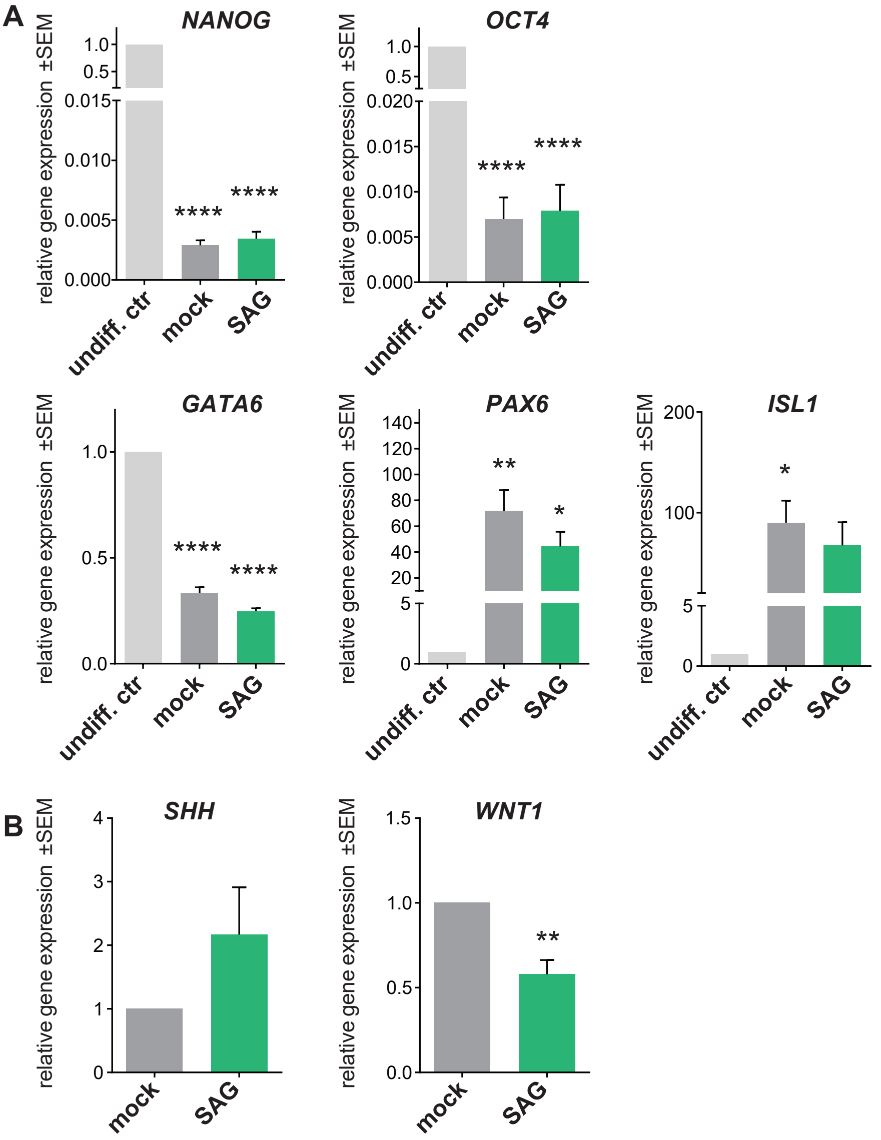


**Supplementary Figure 1.**
**A)** mRNA expression (qRT-PCR) of selected markers in mock- and SAG-treated CCTL14 rosettes on D9; n=5, one-way ANOVA with Tukey's multiple comparisons test. **B)** mRNA expression (qRT-PCR) of selected markers in mock- and SAG-treated CCTL14 rosettes on D20; n=2, t-test.
