## Supplementary Figure 2 for "Tau tubulin kinase 1 and 2 regulate ciliogenesis and human pluripotent stem cells–derived neural rosettes"

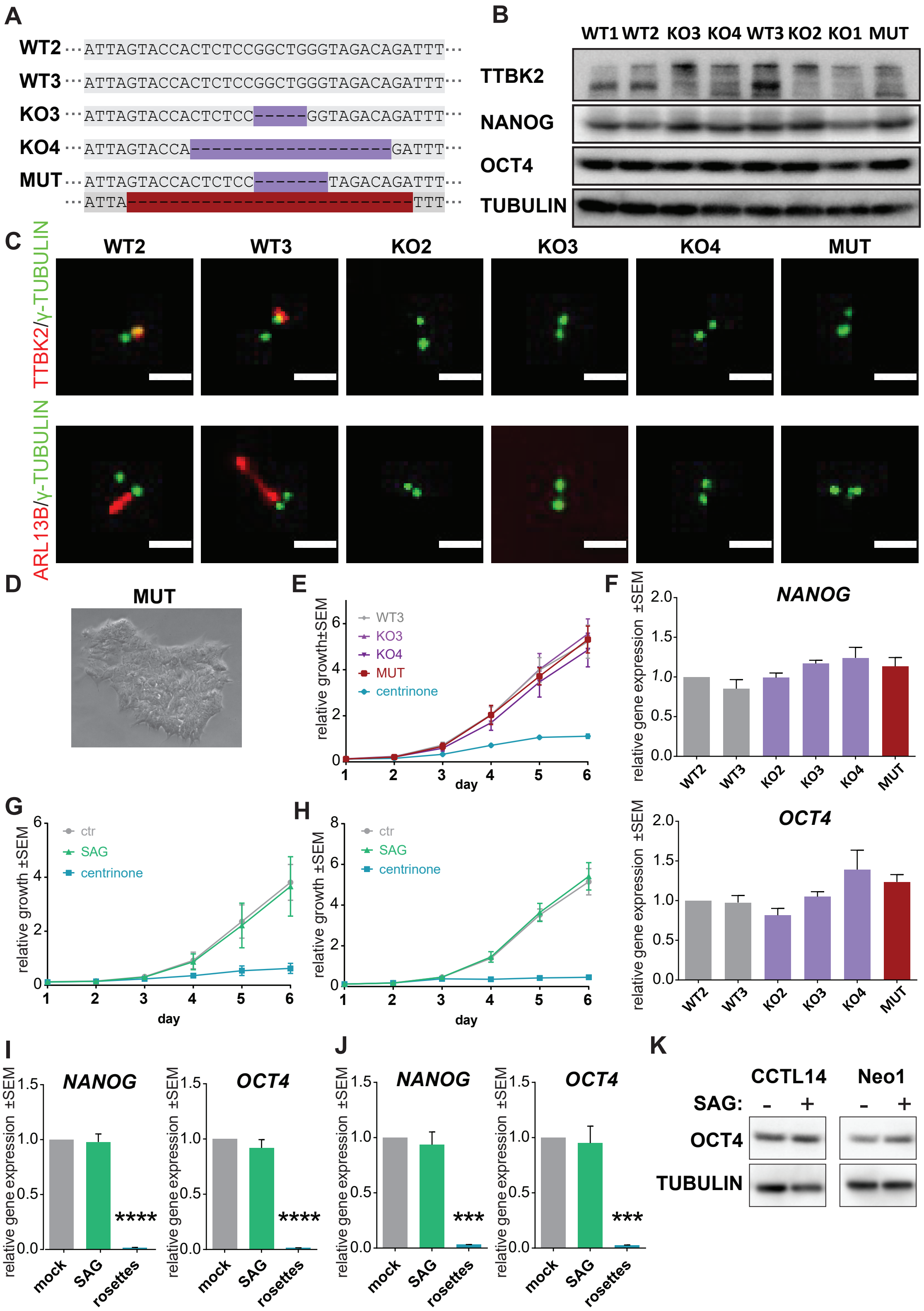


**Supplementary Figure 2.**

**A)** Schematic of the TTBK2 exon 4 sequence detail in WT and TTBK2 KO/MUT cell lines, purple=insertion/deletion, dark red=in frame deletion (mutant). **B)** Representative images of western blot detection of TTBK2, NANOG and OCT4 protein expression in undifferentiated WT and TTBK2 KO/MUT lines; α-TUBULIN was used as a loading control. **C)** Representative images of IF detection of TTBK2 (top) and primary cilia (bottom; visualized by ARL13B staining) in undifferentiated WT and TTBK2 KO/MUT lines, γ-TUBULIN staining was used to detect centrioles; scalebar=2μm. **D)** Representative image of colony morphology of undifferentiated cells in TTBK2 MUT line. **E)** Relative growth comparison of indicated undifferentiated WT and TTBK2 KO/MUT lines assessed by crystal violet absorption measurement, centrinone treatment previously shown to impair the proliferation capacity was used as a control; n=3. **F)** mRNA expression (qRT-PCR) of pluripotency markers NANOG and OCT4 in indicated undifferentiated WT and TTBK2 KO/MUT lines; n=4, one-way ANOVA with Holm-Sidak's multiple comparisons test. **G)** Relative growth comparison of mock- and SAG-treated WT CCTL14 line assessed by crystal violet absorption measurement, centrinone treatment previously shown to impair the proliferation capacity was used as a control; n=3 **H)** Relative growth comparison of mock- and SAG-treated WT Neo1 line assessed by crystal violet absorption measurement, centrinone treatment previously shown to impair the proliferation capacity was used as a control; n=3. **I)** mRNA expression (qRT-PCR) of pluripotency markers NANOG and OCT4 in mock- and SAG-treated (48h) WT undifferentiated CCTL14 line, CCTL14 differentiated into neural rosettes was included for reference; n=3, one-way ANOVA with Tukey's multiple comparisons test **J)** mRNA expression (qRT-PCR) of pluripotency markers NANOG and OCT4 in mock- and SAG-treated (48h) WT undifferentiated Neo1 line, CCTL14 differentiated into neural rosettes was included for reference; n=3, one-way ANOVA with Tukey's multiple comparisons test. **K)** Representative images of western blot detection of pluripotency marker OCT4 in mock- and SAG-treated (48h) WT undifferentiated CCTL14 and Neo1; α-TUBULIN was used as a loading control.
