## Supplementary Figure 3 for "Tau tubulin kinase 1 and 2 regulate ciliogenesis and human pluripotent stem cells–derived neural rosettes"

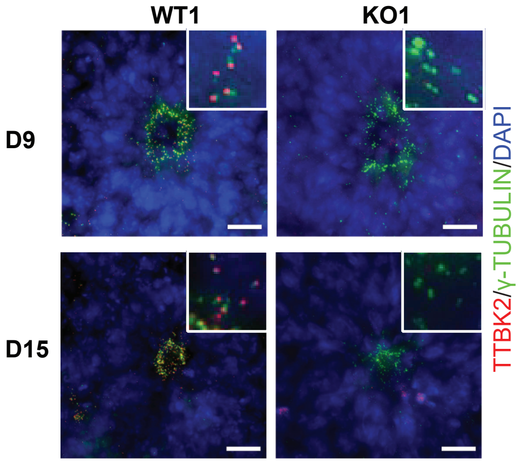


**Supplementary Figure 3.**
Representative images of IF detection of TTBK2 presence in WT1- and TTBK2 KO1-derived neural rosettes on D9 (top) and D15 (bottom), γ-TUBULIN was used to detect centrioles; scalebar=20μm.
