## Supplementary Figure 4 for "Tau tubulin kinase 1 and 2 regulate ciliogenesis and human pluripotent stem cells–derived neural rosettes"

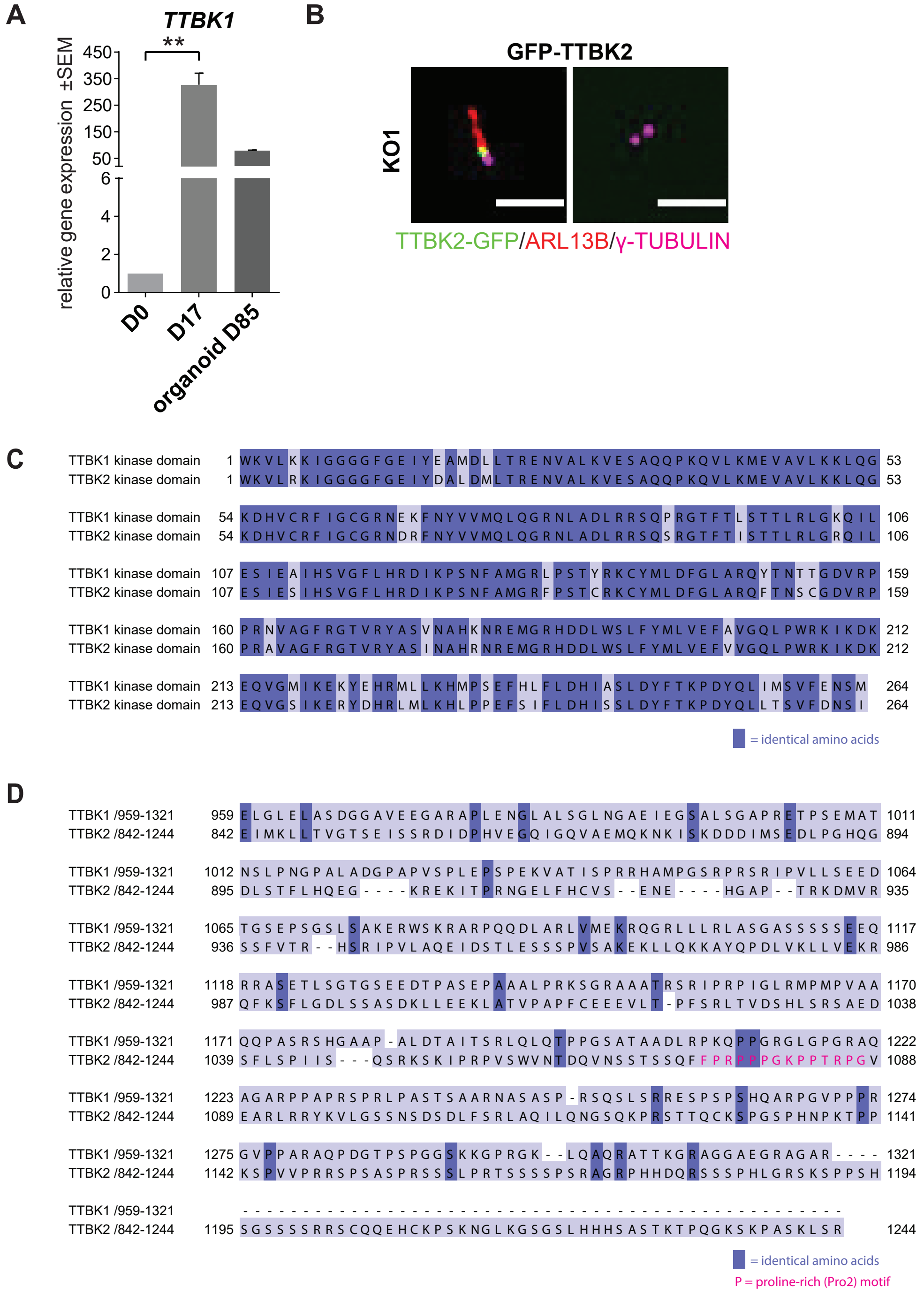


**Supplementary Figure 4.**
**A)** mRNA expression (qRT-PCR) of TTBK1 in undifferentiated parental cell line i3N compared to differentiated neurons on D17 and organoid on D85; n=2, one-way ANOVA with Holm-Sidak's multiple comparisons test. **B)** Representative images of IF detection of primary cilia (visualized by ARL13B staining) in TTBK2 KO1 cells transfected with GFP-TTBK2 (left) or not transfected (right), γ-TUBULIN was used to detect centrioles, scalebar=3μm. **C)** N-terminal TTBK1 and TTBK2 kinase domains alignment, identical amino acids are shown in dark blue. **D)** C-terminal CEP164-binding region in TTBK2 (amino acids 842-1244) aligned to TTBK1, identical amino acids are shown in dark blue, Proline-rich motif necessary for CEP164 binding is highlighted in magenta.
